# Dark accumulation of downstream glycolytic intermediates confers robust initiation of photosynthesis in cyanobacteria

**DOI:** 10.1101/2022.04.04.486922

**Authors:** Kenya Tanaka, Tomokazu Shirai, Christopher J. Vavricka, Mami Matsuda, Akihiko Kondo, Tomohisa Hasunuma

## Abstract

Photosynthesis must maintain stability and robustness throughout fluctuating natural environments. In cyanobacteria, dark-to-light transition leads to drastic metabolic changes from dark respiratory metabolism to CO_2_ fixation through the Calvin-Benson-Bassham (CBB) cycle using energy and redox equivalents provided by photosynthetic electron transfer. Previous studies showed that catabolic metabolism supports the smooth transition into CBB cycle metabolism. However, metabolic mechanisms for robust initiation of photosynthesis are poorly understood due to lack of dynamic metabolic characterizations of dark-to-light transitions. Here, we show rapid (on a time scale of seconds) dynamic changes in absolute metabolite concentrations and ^13^C tracer incorporation after strong or weak light irradiation in the cyanobacterium *Synechocystis* sp. PCC 6803. Integration of this data enables estimation of time-resolved nonstationary metabolic flux underlying CBB cycle activation. This dynamic metabolic analysis indicates that downstream glycolytic intermediates including phosphoglycerate and phosphoenolpyruvate accumulate under dark conditions as major substrates for initial CO_2_ fixation. Compared with wild-type *Synechocystis*, significant delays in the initiation of oxygen evolution are observed in 12 h dark preincubated mutants deficient in glycogen degradation or oxidative pentose phosphate pathway (*Δzwf, Δgnd*, and *ΔglgP*). Accordingly, the degree of delay in the oxygen evolution initiation is proportional to the accumulated pool size of the glycolytic intermediates. These observations indicate that the accumulation of glycolytic intermediates is essential for efficient metabolism switching under fluctuating light environments.

## Main text

Photosynthetic organisms have evolved various mechanisms to maintain robust photosynthesis processes throughout natural environmental perturbations. Ancestral oxygenic and phototrophic cyanobacteria contribute to global carbon cycles by photosynthetic CO_2_ fixation through the Calvin-Benson-Bassham (CBB) cycle^1–3^. The driving forces of the CBB cycle are ATP and NADPH produced by light energy conversion in photosynthetic electron transfer (PET) reactions; and cyanobacteria have evolved metabolic control mechanisms to coordinate CBB cycle activity with environmental light-dark conditions. One mechanism is enzymatic activity regulation that responds to light-dependent cellular environmental changes. For example, glyceraldehyde 3-phosphate dehydrogenase (GAPDH) and phosphoribulokinase (PRK) are regulated by ternary complex formation with the regulatory protein CP12, depending on redox states of specific cysteine residues and cellular NAD/NADP ratio^4–6^.

Maintaining optimal metabolite concentrations is another important mechanism of CBB cycle regulation. In general, concentration robustness is necessary for maintenance of cellular functionality^7^. Recent analyses performed using a theoretical photosynthetic model have predicted that maintaining the appropriate concentrations of CBB cycle intermediates is crucial for the initiation of CO_2_ fixation^8^. Experimental evidence in cyanobacteria has confirmed that the transfer of the carbon supply from a storage polymer (glycogen) to the CBB cycle affects photosynthetic efficiency^9,10^. In addition, while enzymatic regulation systems control metabolism, metabolite concentration affects metabolism itself by thermodynamic constraints and effects on enzymatic kinetics as represented by the Michaelis-Menten equation^11,12^. Thus proper regulation of metabolite distribution is considered to be important for activation of the CBB cycle.

However, the metabolic distribution underlying robust CBB cycle operation under light-dark perturbations is still unclear. Ascertaining which metabolite is important for the CBB cycle kickstart requires understanding metabolic dynamics during CBB cycle activation immediately after a dark-to-light transition. For this, we aim to derive dynamic metabolic flux changes during photosynthetic activation. Although metabolic flux analysis (MFA) using an isotope tracer enables the mapping of intracellular carbon fluxes^13–15^, the approach cannot be applied to a metabolically non-stationary state with variable metabolic fluxes and pool sizes. On the other hand, previously developed dynamic MFA, which incorporates specialized statistical approaches, has permitted the estimation of long-term flux changes over the culture interval^16,17^. There has been (to our knowledge) no appropriate approach for the quantitative determination of metabolic flux changes throughout short, metabolically transient states, including that of CBB cycle activation.

In this study, we successfully quantify absolute metabolite concentration changes at second-scale intervals after dark-to-light transitions in the cyanobacterium *Synechocystis* sp. PCC 6803 (hereafter *Synechocystis*). Additionally, to identify metabolic pathways that are activated during CBB cycle initiation, dynamic ^13^C labeling is performed immediately following light irradiation. Accurate concentration and labeling dynamics allow us to conduct short-time dynamic metabolic flux analysis (STD-MFA) to estimate non-stationary flux changes during activation of the CBB cycle. The STD-MFA results show that highly accumulated glycolytic intermediates including 3-phosphoglycerate (3PG), 2-phosphoglycerate (2PG), and phosphoenolpyruvate (PEP) are main substrates for initial CO_2_ fixation. This metabolic phenomenon is further characterized by the delay of oxygen evolution in several mutant strains, in which the accumulated pool sizes of the glycolytic intermediates are found to be smaller than that of wild-type cells.

## Results

### Strategy for estimating metabolic flux dynamics

To estimate non-stationary metabolic flux changes, the following relationship is employed. The accumulation rate of a metabolite in a metabolic network is defined as the sum of all fluxes producing the metabolite minus the sum of all fluxes consuming the metabolite, as represented by following equation:

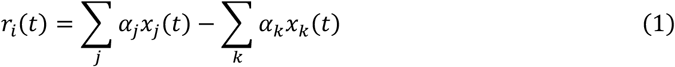

where, *x*_*j*_*(t)* is the flux through reaction *j, α*_*j*_ is a stoichiometric coefficient, and *r*_*i*_*(t)* is the accumulation rate of metabolite *i*^18^. An example with three metabolic fluxes is illustrated in Fig. 1a. To obtain accumulation rates for estimation of the flux values, it is necessary to examine the absolute concentration changes of metabolites with high time resolution. Namely, the metabolic flux during CBB cycle activation can be calculated by quantifying the changes in metabolite concentrations over highly time-resolved intervals immediately following light irradiation.

**Fig. 1.**
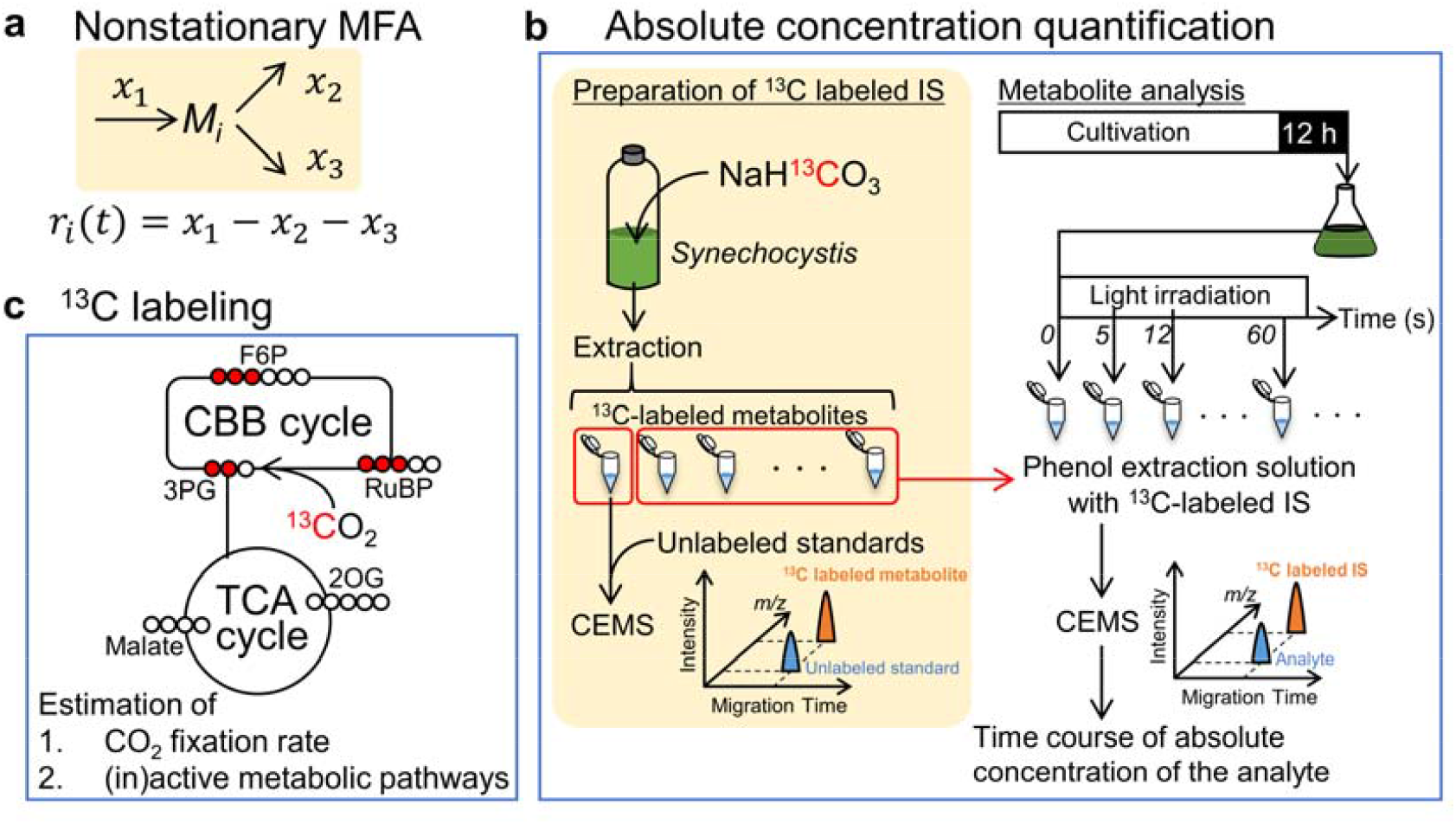
Process for estimating temporal changes in metabolic flux. **a**, Relationship between concentration change and metabolic flux. *r*_*i*_(t) is the accumulation rate of metabolite *M*_*i*_. *x*_*n*_ represents the metabolic flux of the *n*th reaction. **b**, Experimental procedure for quantifying absolute metabolite concentration. **c**, Schematic example of isotope-labeled state during ^13^CO_2_ incorporation. Empty and red-filled circles represent unlabeled and labeled carbon atoms, respectively. Abbreviations: CBB, Calvin-Benson-Bassham; CEMS, capillary electrophoresis-mass spectrometry; F6P, fructose 6-phosphate; IS, internal standard; RuBP, ribulose 1,5-bisphosphate; TCA, tricarboxylic acid; 2OG, 2-oxoglutarate; 3PG, 3-phosphoglycerate.

### Absolute concentration changes in central carbon metabolism

We sought to determine the absolute concentration dynamics of central energy metabolites in *Synechocystis*. As light irradiation rapidly causes metabolic variations driven by increases in ATP and NADPH via the PET reaction, rapid sampling and quenching of cellular metabolic activity are necessary to accurately quantify metabolic changes during CBB cycle activation. A standardized procedure for intracellular metabolome quantification requires multiple analytical steps, including quenching of microbial metabolic activity, separation of microbial cells from growth medium, extraction of intracellular metabolites, and finally mass spectrometric metabolite analysis. These multistep processes largely impede accurate estimation of the absolute ‘*in vivo*’ cellular metabolic state^19–21^.

To overcome the above problems of the present study, a combination of two strategies were employed. First, to analyze intracellular metabolites, cells are separated from the growth medium and suspended in 1 mM NaHCO_3_ prior to metabolite sampling to remove extracellular metabolites and medium components. Note that the exchange of medium for NaHCO_3_ solution is a typical pretreatment for measuring photosynthetic activity of fresh-water cyanobacteria^22^. Carefully dropping the cell suspension into the extraction solution during light irradiation enables the rapid quenching and extraction of intracellular metabolites and avoids metabolic state variations during sampling. A phenol-chloroform-isoamyl alcohol (PCI) solution (pH 7.9) is selected as the extraction solution because PCI extraction enables rapid and efficient extraction of NADPH from *Synechocystis*^23^. Second, stable isotopomers of the analyte compounds are used as internal standards (IS). Experimental protocols using isotope-labeled IS are available for absolute quantification of metabolite concentrations in various microorganisms including cyanobacteria^24–26^.

Figure 1b shows the experimental process for the exact determination of absolute metabolite concentration dynamics. A ^13^C-labeled IS mixture is prepared by extracting metabolites from *Synechocystis* cells cultivated with NaH^13^CO_3_ as the sole carbon source. Absolute concentrations of uniformly ^13^C-labeled metabolites are determined using capillary electrophoresis-mass spectrometry (CE-MS) based on the peak area ratios of unlabeled metabolites (with known concentration) and uniformly ^13^C-labeled metabolites. Then, to quantify absolute concentration changes during CBB cycle activation, dark-adapted *Synechocystis* cells (grown in darkness for 12 h) are illuminated at a light intensity of 30 μmol m^−2^ s^−1^ (same as the growth light intensity) or 200 μmol m^−2^ s^−1^. Metabolites of these cells are rapidly extracted using PCI solvent containing the ^13^C-labeled extract as labeled IS with concentrations determined from another aliquot of the same extraction batch (Fig. 1b). Because of high physicochemical similarities between labeled and unlabeled central metabolites, both isotopomers undergo similar processes of degradation and ion suppression, and therefore accurate concentrations can be calculated^24–26^.

Given that absolute metabolite concentrations are determined based on the ratio between unlabeled and uniformly labeled peak areas (Fig. 1b), the process is expected to prevent contamination of unlabeled metabolites in the ^13^C-labeled IS. The proportions of uniformly ^13^C-labeled metabolites (compared to the sum of unlabeled and uniformly labeled metabolites) are found to exceed 95% for all metabolites except for citrate (Supplementary Fig. 1). Figure 2 shows temporal changes in absolute metabolite concentrations after the initiation of light irradiation at 30 μmol m^−2^ s^−1^ or 200 μmol m^−2^s^−1^. It should be noted that a small amount of some metabolites are extracellular (Supplementary Fig. 2). A previous study that measured *in vivo* NADPH fluorescence revealed that the amount of NADPH in *Synechocystis* increases within 0.5 s of illumination^27^. The rapid increase of NADPH observed within 5 s in the present work is consistent with that of the earlier *in vivo* study^27^, confirming that the current metabolomics strategy can successfully detect the rapid start of photosynthesis. As with NADPH, the level of the energy cofactor ATP also increases rapidly. Overall, the levels of 3PG, 2PG, and PEP decrease; the levels of other CBB cycle intermediates (except 3PG, ribulose 5-phosphate (Ru5P) and xylulose 5-phosphate (Xu5P)) increase, while the levels of tricarboxylic acid (TCA) cycle intermediates do not change appreciably. Considering these metabolite variations together, we inferred that accumulated glycolysis intermediates including 3PG, 2PG, and PEP likely were converted to CBB cycle intermediates at the photosynthetic start.

**Fig. 2.**
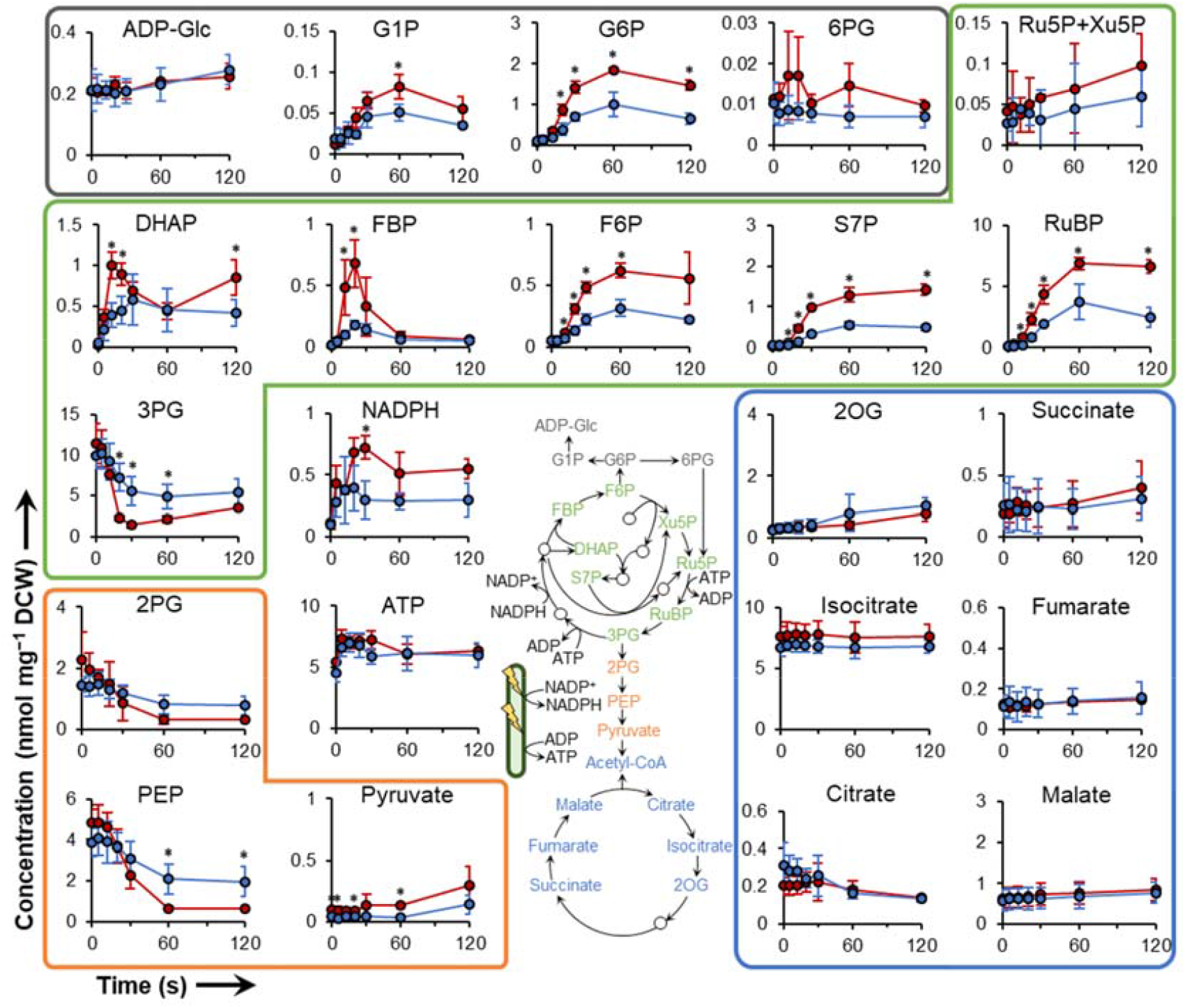
Rapid temporal changes in absolute metabolite concentrations during the initiation of photosynthesis. Intracellular metabolites are extracted at the indicated times after transition from darkness to light-irradiated condition at light intensities of 200 μmol m^−2^ s^−1^ (red line) or 30 μmol m^−2^ s^−1^ (blue line). Levels of glyceraldehyde 3-phosphate (GAP), erythrose 4-phosphate (E4P), and ribose 5-phosphate (R5P) are below the detection limit (0.001 nmol/mg DCW). 1,3-Bisphosphoglycerate (BPG) and sedoheptulose 1,7-bisphosphate (SBP) are not identified due to lack of commercial reagent for use as a standard. Metabolites are grouped by different colors according to the central schematic metabolic network. Abbreviations: DCW, dry cell weight; DHAP, dihydroxyacetone phosphate; FBP, fructose 1,6-bisphosphate; F6P, fructose 6-phosphate; G1P, glucose 1-phosphate; G6P, glucose 6-phosphate; 2OG, 2-oxoglutarate; PEP, phosphoenolpyruvate; 6PG, 6-phosphogluconate; 2PG, 2-phosphoglycerate; 3PG, 3-phosphoglycerate; RuBP, ribulose 1,5-bisphosphate; Ru5P, ribulose 5-phosphate; S7P, sedoheptulose 7-phosphate; Xu5P, Xylulose 5-phosphate. Values are the mean ± SD (bars) of three independent experiments. Significant differences between the two light intensities were evaluated by a two-tailed non-paired Student’s t test (* *p* < 0.05).

### ^13^C labeling kinetics

A direct way to confirm this hypothesis is to determine the metabolic flux distribution. However, precise metabolic flux values cannot be determined when the number of unknown flux parameters is larger than the number of known metabolite concentration changes in the metabolic flux model. To accurately perform STD-MFA, we sought to refine our metabolic models by excluding metabolic pathways that are inactive during photosynthetic activation. The metabolite concentration data shown in Fig. 2 cannot identify inactive metabolic pathways. Even if the metabolic concentration change is zero, metabolic flux is not necessarily zero, because such an observation instead may indicate that metabolic inflow and outflow are balanced. In the case where the concentration of a metabolite is stable and ^13^C is not incorporated into the metabolite during photosynthetic ^13^CO_2_ fixation, the metabolic fluxes around the metabolite are estimated to be very small, and therefore can be neglected. Accordingly, we next measured the dynamic ^13^C isotope incorporation after the transition from dark to light conditions, by starting photosynthesis while the cells are suspended in a solution of NaH^13^CO_3_ (Fig. 1c). As a result, a transient increase in ^13^C fractions of CBB cycle intermediates is observed (Fig. 3a, Supplementary Fig. 3). On the other hand, labeling of 6-phosphogluconate (6PG) and TCA cycle intermediates (except for malate and fumarate) is not observed. In addition, a low ^13^C fraction of the glycogen precursor ADP-glucose^28^, is observed. When considered together with the stable concentrations of TCA and oxidative pentose phosphate (OPP) pathway intermediates (Fig. 2), these results suggest extremely low metabolic fluxes of the TCA cycle, OPP pathway, Entner-Doudoroff (ED) pathway, and glycogen synthesis pathway, immediately after light irradiation.

**Fig. 3.**
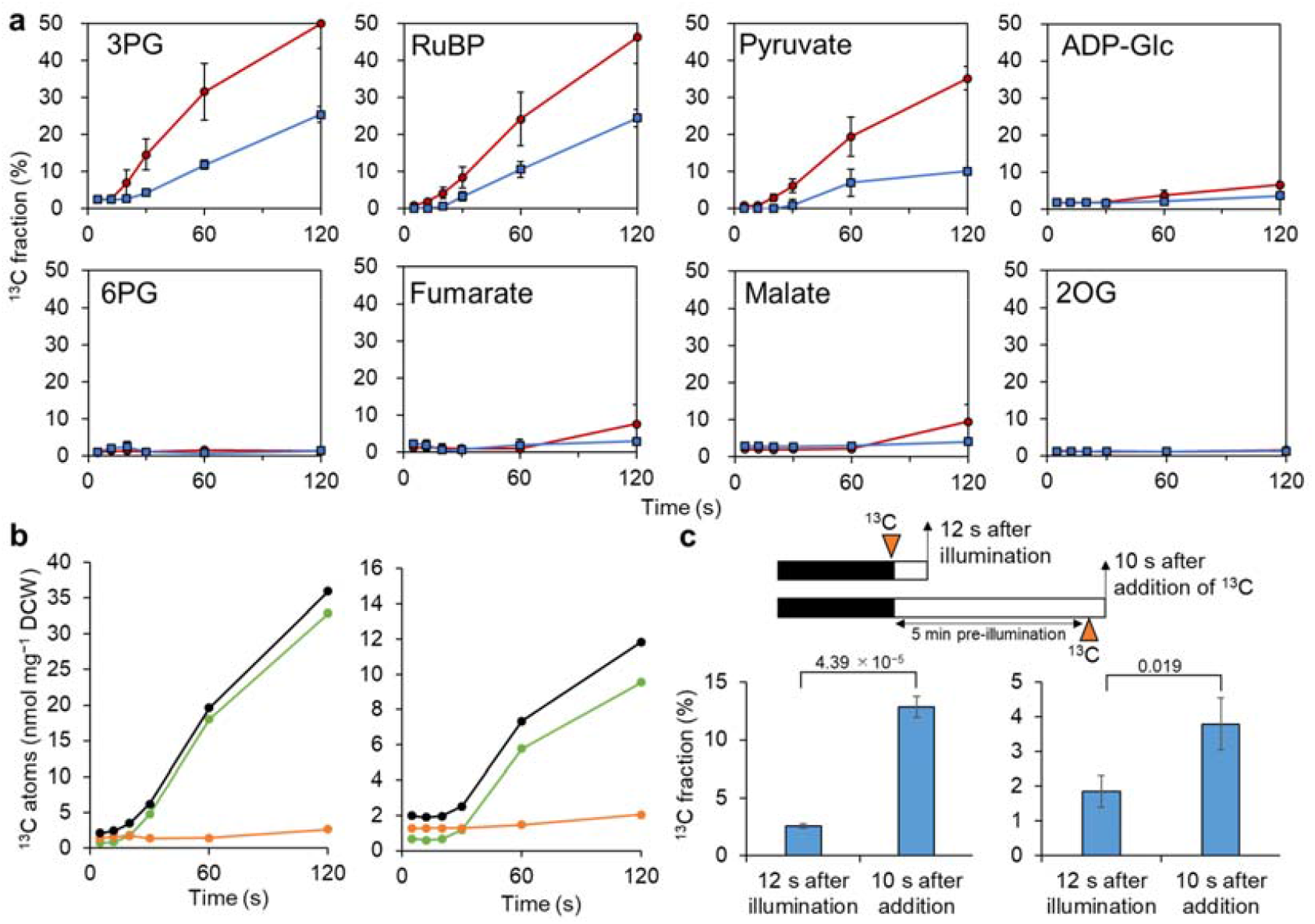
Incorporation of ^13^C atoms during CBB cycle activation. (a) Time course of ^13^C fraction of metabolites following light irradiation at 200 μmol m^−2^ s^−1^ (red line) or 30 μmol m^−2^ s^−1^ (blue line). Values are the mean ± SD (bars) of three biological replicates. (b) Total absolute amount of ^13^C-labeled carbon atoms. The number of incorporated ^13^C atoms is derived from absolute concentration and ^13^C fraction data. Black, green, and orange lines indicate total amount of ^13^C atoms of all measured metabolites, CBB cycle and OPP pathway metabolites (G6P, 6PG, and metabolites in green zone in Fig. 2), and TCA cycle and lower glycolysis metabolites (2PG, PEP, pyruvate, and metabolites in yellow zone in Fig. 2). Left panel: 200 μmol m^−2^ s^−1^, right panel: 30 μmol m^−2^ s^−1^. (c) Effect of pre-illumination on the start of ^13^C incorporation. ^13^C fractions of 3-phosphoglycerate (left panel) and ribulose 1,5-bisphosphate (right panel) after 12 s of illumination (200 μmol m^−2^ s^−1^) of the 12-h dark-adapted cells were lower than those at 10 s after addition of 1 mM NaH^13^CO_3_ to cells that had been pre-illuminated (200 μmol m^−2^ s^−1^) for 5 min. Presumably, the higher ^13^C incorporation rate in pre-illuminated cells reflects the fact that CO_2_ fixation already had been activated (by pre-illumination) prior to the addition of NaH^13^CO_3_. Thus, lower ^13^C incorporation by dark-adapted cells can be attributed to the low activation state of CO_2_ fixation during photosynthetic induction. Values are the mean ± SD (bars) of three biological replicates. Significant differences were evaluated by a two-tailed non-paired Student’s t test; *p* values were as indicated.

In addition to ribulose bisphosphate carboxylase/oxygenase (RuBisCO), the primary metabolic enzymes that incorporate CO_2_ are phosphoenolpyruvate carboxylase (PepC) and malic enzyme (ME), which are both related to malate metabolism. Since malate is not labeled until 60 s after the initiation of light irradiation (at both of the tested light intensities), ^13^C incorporation before 60 s is inferred to reflect RuBisCO activity. The amount of incorporated ^13^C is calculated by multiplying the measured ^13^C fraction (Fig. 3a, Supplementary Fig. 3) by the absolute carbon concentration of each metabolite (shown in Fig. 2); these analyses are presented in Supplementary Fig. 4. The total amount of incorporated ^13^C is derived by summing the number of ^13^C atoms of all measured metabolites (Fig. 3b). Notably, ^13^C incorporation does not start until 12 s (200 μmol m^−2^ s^−1^) or 20 s (30 μmol m^−2^ s^−1^) after the initiation of light irradiation. On the other hand, ^13^C incorporation is observed at least 10 s after the addition of NaH^13^CO_3_ to cells with activated photosynthesis in response to 5 min of illumination (Fig. 3c). Together, these results suggest that the observed delay in the start of ^13^C incorporation is not due to cellular uptake of NaH^13^CO_3_. Instead, these data indicate that RuBisCO-mediated CO_2_ fixation remains inactive during the initial stage of photosynthetic metabolism, immediately following the start of illumination. This delayed onset of CO_2_ fixation is consistent with the photosynthetic induction phenomena observed in other photosynthetic organisms^29^.

It is interesting to note that labeled carbon is incorporated into malate and fumarate after 120 s, particularly in cells illuminated at 200 μmol m^−2^ s^−1^, at a time when other TCA cycle intermediates remained unlabeled (Supplementary Fig. 3). This result suggests that malate and fumarate are generated by the reductive TCA pathway, which is known as a fermentation pathway dependent on NADH oxidation^30^. Thus, the accumulation of NADH may drive the activity of this reductive TCA pathway (Supplementary Fig. 5).

### Short-term dynamic metabolic flux analysis (STD-MFA)

Temporal changes in net metabolic fluxes are calculated based on the obtained data. As mentioned above, during the first 60 s of illumination, the observed net metabolic fluxes in pathways other than the CBB cycle and glycolysis are very small. In fact, the concentrations of newly synthesized amino acids do not change significantly within 60 s, even in cells illuminated at 200 μmol m^−2^ s^−1^, although the labeled levels of some amino acids (serine, alanine, phenylalanine, tyrosine and ornithine) increased significantly after the first minute of illumination (Supplementary Fig. 6). Furthermore, photorespiratory intermediates such as glycolate, glyoxylate and glycerate do not accumulate during the first 60 s of illumination (Supplementary Fig. 7). Therefore, the metabolic pathways shown in Supplementary Fig. 8a are used for subsequent calculations, focusing on metabolic dynamics during the first 60 s following the initiation of illumination.

To solve equation (1) and conduct STD-MFA, accumulation rates for metabolites in the metabolic model shown in Supplementary Fig. 8a are derived from the slopes obtained between two consecutive concentration data points in Fig. 2. In addition to our use of equation (1), the CO_2_ fixation rate is calculated as the increase in the rate of total ^13^C incorporation (Fig. 3b). Specifically, the latter parameter is interpreted as the metabolic flux from ribulose 1,5-bisphosphate (RuBP) to 3PG, given that the ^13^C incorporation experiment mentioned above indicates that all carbon atoms fixed during the first 60 s of illumination are attributable to RuBisCO activity. The net metabolic fluxes of each time interval are estimated by solving these equations simultaneously (Fig. 4, Supplementary Fig. 8). The estimated flux values are validated and judged to be reliable, given that almost all of the estimated metabolite accumulation rate values (which were derived from the estimated fluxes) yielded values that are within the mean ± SD range of the measured accumulated rate (Supplementary Fig. 9). For more details of the calculation process and equations, see the Methods section.

**Fig. 4.**
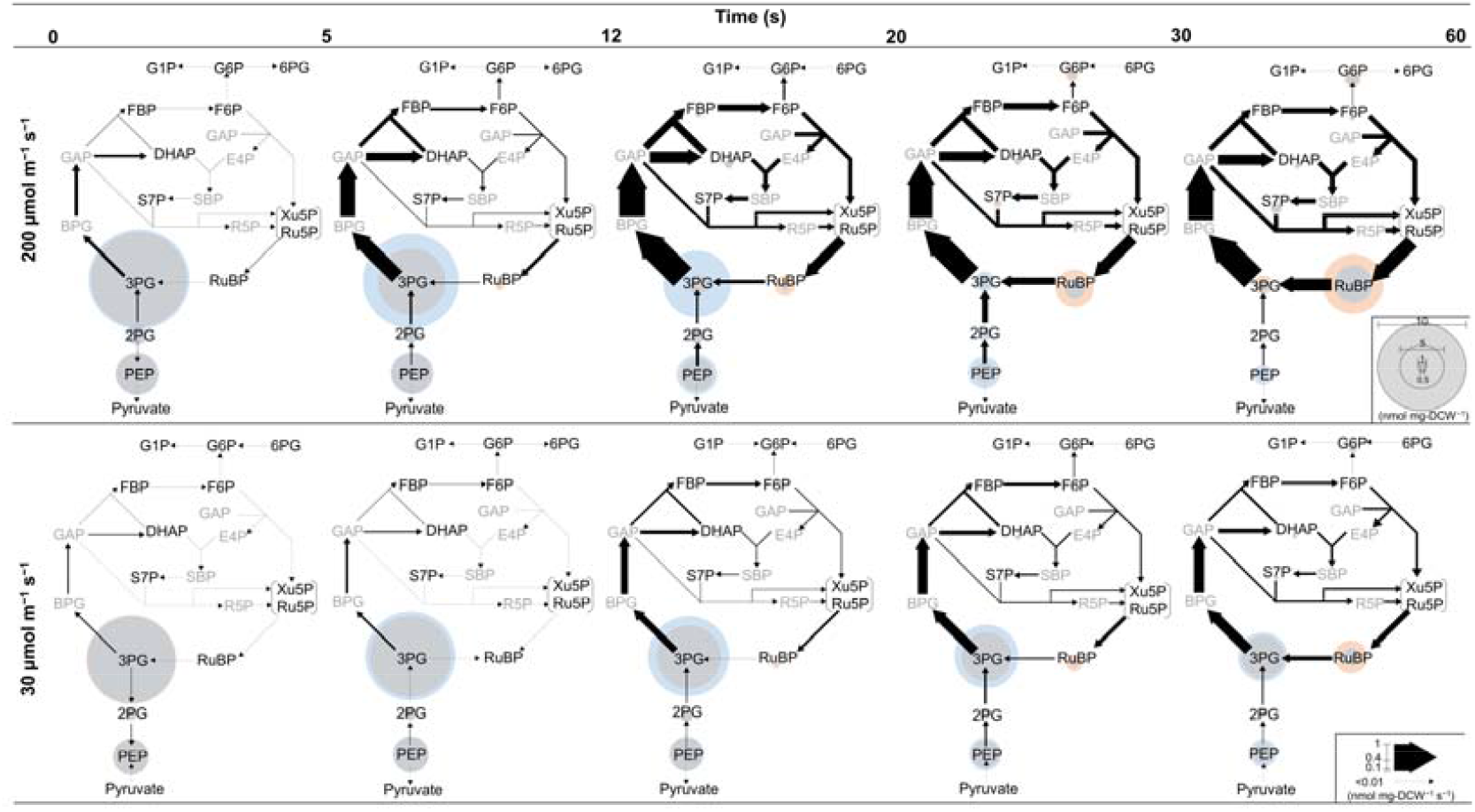
Estimated change in metabolic flux after initiation of light irradiation at 200 μmol m^−2^ s^−1^ (upper row) and 30 μmol m^−2^ s^−1^ (lower row). Metabolic fluxes are indicated by the thickness of arrows with values indicated by the scale bar in the lower right corner. Metabolite concentrations at the beginning and end of each time interval are indicated by the diameters of blue and orange circles, respectively. Overlap between orange and blue circles (namely invariant concentration during a time interval) is shown in gray. Abbreviations are the same as in the Figure 2 legend.

The time course of metabolic flux and metabolite concentration distribution in Fig. 4 clearly indicates that photosynthetic metabolism in *Synechocystis* starts by converting dark accumulated 3PG as an initial substrate into RuBP under both two light intensities. On the other hand, the reduction fluxes of 3PG differ greatly between cells illuminated at 30 and 200 μmol m^−2^ s^−1^, even before the initiation of CO_2_ fixation (i.e. during the first 5-20 s). In steady-state photosynthesis, the increased rate of electron donor (NADPH) production from the PET reaction at the higher light intensities is balanced by the rate of electron acceptor (3PG) production that is increased by the elevated CO_2_ fixation rate. On the other hand, before the onset of CO_2_ fixation that follows the initiation of light irradiation, excess reducing power can be accumulated because 3PG cannot be produced while NADPH is produced by the PET reaction. Nevertheless, the increase in the reducing fluxes of 3PG with the higher light intensity (200 μmol m^−2^ s^−1^) before CO_2_ fixation suggests that, even before the onset of CO_2_ fixation, the rate of 3PG consumption is balanced by the rate of NADPH production from the PET reaction, which varies with light intensity. In addition, it can clearly be seen that 2PG and PEP are converted to 3PG under both light intensities (Fig. 4). Therefore, we propose that the accumulation of these glycolytic intermediates provides a “stand-by” state which precedes robust initiation of photosynthesis; thus, these intermediates are expected to play an important role in balancing cofactor regeneration and consumption during activation of the CBB cycle.

### Correlation between dark-accumulation of glycolytic intermediates and oxygen evolution

The effects of glycolytic intermediate accumulation on the initiation of photosynthesis are further examined using mutants lacking enzymes involved in dark metabolism. Under dark conditions, cyanobacteria obtain NADPH mainly via the OPP pathway^31–33^. As an initial OPP pathway substrate, glucose 6-phosphate (G6P) is supplied by glycogen degradation through glycogen phosphorylase. In a glycogen phosphorylase-deficient mutant of *Synechocystis*, decreased accumulation of CBB cycle metabolites has been suggested as a cause of delayed oxygen evolution after dark incubation^9^. However, the key metabolites that determine CBB cycle activation efficiency have not been reported.

To better understand CBB cycle activation, the relationship between absolute metabolite concentration and oxygen evolution activity, following 12 h dark preincubation periods, is characterized in mutants lacking glycogen phosphorylase, glucose 6-phosphate dehydrogenase or 6-phosphogluconate dehydrogenase (*ΔglgP, Δzwf, Δgnd*). In all strains, the increase of oxygen evolution rate (OER) is significantly delayed when following a dark preincubation period (Fig. 5a). Especially, the OER of dark preincubated *Δgnd* does not increase during 5 min of light irradiation. To quantify OER activation efficiency, ratios of OER values at 60 s after light irradiation of dark preincubated cells to that of non-dark incubated cells are calculated (Fig. 5b). The resulting OER ratio is highest in wild-type (WT) *Synechocystis*, followed by *Δzwf, ΔglgP, Δgnd*, in descending order.

**Fig. 5.**
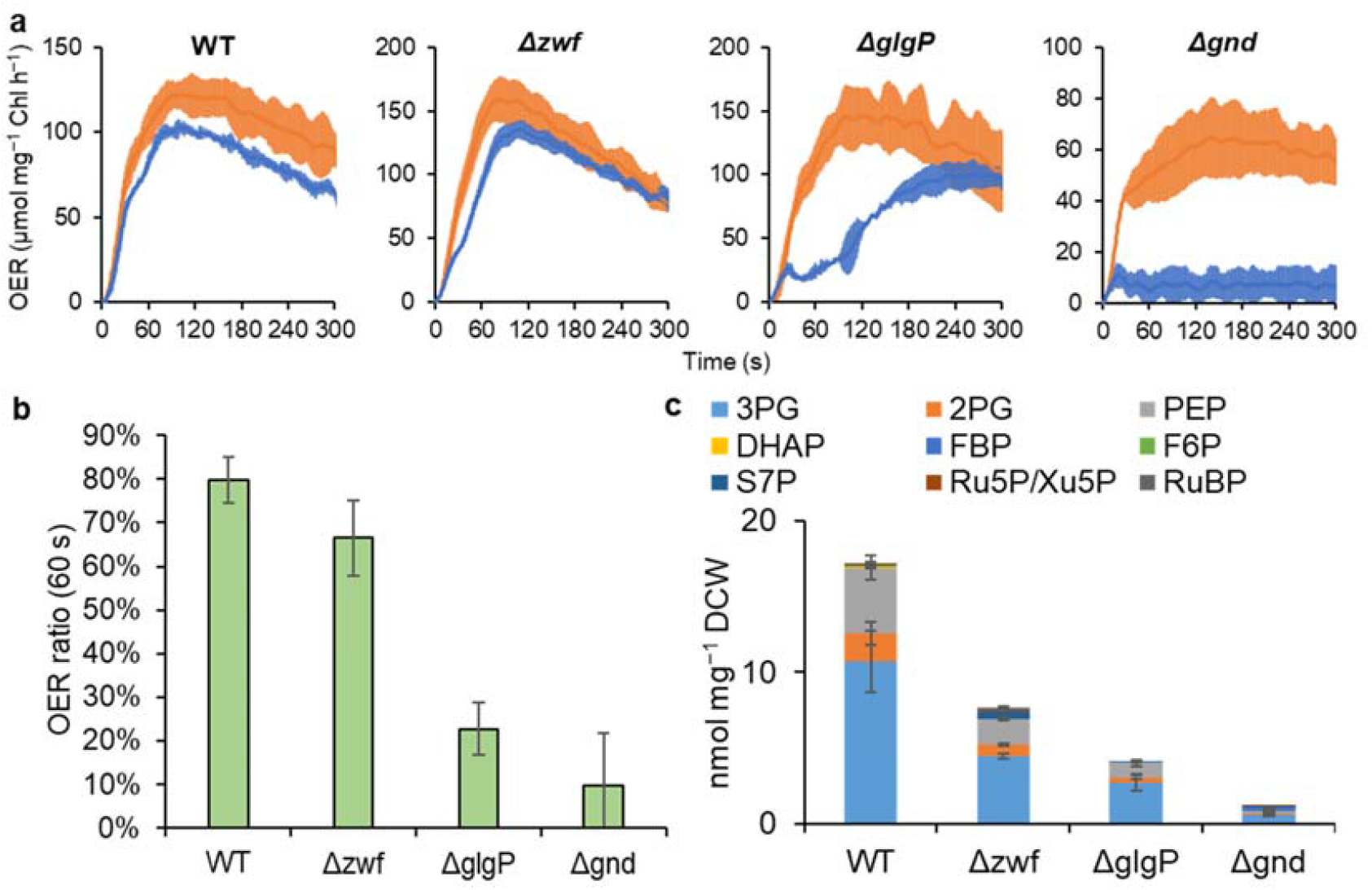
Effect of dark metabolism on initiation of photosynthesis and accumulation of substrates for initial CO_2_ fixation. (a) Oxygen evolution rate (OER) of wild-type (WT) and mutants. Orange and blue lines indicate OER transients of light-adapted and 12 h-dark-adapted cells after light irradiation (200 μmol m^−2^ s^−1^). (b) Ratio of OER value of dark-adapted cells to that of light-adapted cells at 60 s are compared among mutants. (c) Absolute concentrations of substrates for initial CO_2_ fixation in 12 h dark-adapted cells. Values are the mean ± SD (bars) of three biological replicates. Abbreviations are the same as in the Figure 2 legend.

On the other hand, 9 substrates are found to be used for initial CO_2_ fixation (measured metabolites converted into RuBP during the first 60 s of light irradiation in Fig. 4): 3PG, 2PG, PEP, dihydroxyacetone phosphate (DHAP), fructose 1,6-bisphosphate (FBP), fructose 6-phosphate (F6P), sedoheptulose 7-phosphate (S7P), Xu5P and Ru5P. The total amounts of these initial substrates in the 12 h dark preincubated cells follows the same trend as that of the OER ratios (Fig. 5c). The three glycolytic intermediates (3PG, 2PG, and PEP) account for 98% of the initial substrates in WT, 90% in *Δzwf*, 99% in *ΔglgP*, and 68% in *Δgnd*. Together these results confirm that photosynthesis activation efficiency is clearly correlated with the accumulated amounts of glycolytic intermediates. These results demonstrate that the accumulation of glycolytic intermediates under dark conditions is essential for rapid activation of photosynthetic metabolism.

## Discussion

### A new method for estimating metabolic flux kinetics

The STD-MFA method developed in this study is able to quantify rapid changes in metabolic flux over short intervals (within a few minutes), in response to a dynamic phenomenon. This method requires the measurement of time-resolved absolute concentration changes, uptake or evolution rate of one or some metabolites (in present case, carbon uptake rate) and metabolic model refinement based on isotope incorporation data before flux calculation. Previously developed dynamic metabolic flux analysis methods typically are intended to estimate flux change over long periods of culture processes, and require technically specialized mathematical approaches^16,17^. In contrast, STD-MFA permits the direct determination of metabolic flux changes in exchange, albeit while focusing on short time intervals. We expect that this method also may be of use for understanding other non-stationary rapid metabolic changes that have been the recent focus of attention, such as the end of photosynthesis and adaptation against oxidative stress^34,35^.

### Regulation mechanisms of the CBB cycle

Metabolic regulation of the CBB cycle includes control of enzyme activity and metabolite accumulation. The present study reveals that accumulation of glycolytic intermediates streamlines activation of photosynthetic metabolism. On the other hand, the presence of plant-like enzymatic regulation in cyanobacteria is debatable while plant CBB cycle enzymes are strictly regulated depending upon light-dark conditions^36–40^. The present *in vivo* metabolomic data in Fig. 2 is consistent with the following enzymatic activations in *Synechocystis*.

Metabolic flux of conversion of 3PG remains small until 5 s (200 μmol m^−1^ s^−1^) or 12 s (30 μmol m^−1^ s^−1^), indicating the presence of regulation in phosphoglycerate kinase (PGK) and/or GAPDH. Cyanobacterial PGK and GAPDH are activated by reductive dissociation of cysteine pairs^5,41^. Transient accumulation of FBP indicates the existence of an activation period for the fructose-1,6-/ sedohepturose-1,7-bisphosphatase (FBP/SBPase) enzyme. FBP/SBPase in *Synechocystis* is activated by the reduction of a thiol group and binding of Mg^2+^, and this enzyme is inhibited allosterically by the binding of AMP^42,43^. Indeed, a rapid decrease in AMP concentration was confirmed after light irradiation (Supplementary Fig. 5). In addition, temporal accumulation of DHAP upon irradiation with 200 μmol m^−2^ s^−1^ light may be attributable to the transient rate-limitation of FBP/SBPase. Similar to the behavior of FBP, the accumulation and subsequent decrease of RuBP suggests a phase shift from inactivated to activated RuBisCO. RuBisCO activity is activated by various factors: (a) carbamylation to the activator lysine and binding of a magnesium ion to the carbamate anion as essential regulations^44,45^, (b) effectors such as orthophosphate^46^, several phosphorylated sugars and NADPH^47^, (c) cysteine redox state^48,49^, (d) RuBisCO activase^37^. Among these activation processes, *Synechocystis* lacks RuBisCO activase. In addition, cyanobacteria encapsulate RuBisCO enzymes into protein-encased microcompartments called the carboxysome^50^. Initiation of CO_2_ fixation requires entrance of RuBisCO substrate, RuBP into the carboxysome. Although kinetics of this RuBP uptake is unknown, this process might affect activation kinetics of CBB cycle. Note that similar dynamics of the RuBP pool size has been observed after a dark-to-light transition in plant leaves, and is associated with RuBisCO activation^51,52^.

Free energy analysis provides further insight into the regulation in the CBB cycle. Thermodynamic feasibility is estimated by the reaction Gibbs energy change (Δ_*r*_*G*), calculated based on the following equation:

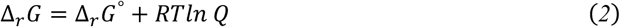

where, Δ_*r*_*G*º is the standard reaction Gibbs energy change, *R* is the ideal gas constant, *T* is the temperature, and *Q* is the reaction quotient, which includes concentration terms. The availability of absolute concentration data enables assessment of the thermodynamically feasible flux directions. Corresponding Δ_*r*_*Gº* values are obtained from the eQuilibrator database^53^ (Supplementary Table 1). Interestingly, conversion of 3PG to DHAP is thermodynamically allowed even under dark conditions as the Δ_*r*_*G* for the reaction from 3PG forming DHAP is a large negative value (Supplementary Fig. 10). This non-equilibrium state supports the presence of dark inactivation of PGK and GAPDH as mentioned above. The Δ_*r*_*G* value of conversion from PEP to pyruvate catalyzed by pyruvate kinase (PK) is found to be a very negative value compared with that of heterotrophic organisms^12^ (Supplementary Fig. 10). Along with GAPDH and PGK, PK activity regulation may contribute to the observed accumulation of 3PG, 2PG, and PEP.

### Essential metabolic pathways for dark accumulation of glycolytic intermediates

Accumulation of glycolytic intermediates decreases in *Δzwf, ΔglgP*, and *Δgnd*, suggesting that the accumulated glycolytic intermediates are derived from glycogen degradation and the OPP pathway. The first step of the OPP pathway is the irreversible conversion of G6P into 6PG by glucose-6-phosphate dehydrogenase (G6PDH). Then, 6PG is decarboxylated into Ru5P by 6-phosphogluconate dehydrogenase (6PGDH). Accumulation of initial CO_2_ fixation substrates in *Δgnd* is much lower than in *Δzwf* (Fig. 5c). This may be explained by the G6PDH-mediated conversion of glycogen-derived G6P into 6PG, with will then accumulate if it is not readily converted to other metabolites. Indeed, 6PG accumulation in *Δgnd* (45.0 nmol mg^−1^ DCW) is about 4000-fold higher than in the wild-type (0.011 nmol mg^−1^ DCW) after dark incubation. On the other hand, G6P in *Δzwf* can be metabolized by other pathways such as the Embden-Meyerhof-Parnas (EMP) pathway, which might prevent the depletion of glycolytic intermediates.

### Significance and conservation of intermediate pool size

Although concentration robustness is generally considered to be a fundamental factor for maintaining cellular functions^7^, the precise role of metabolic robustness in photosynthetic organisms has not been clarified. Balancing photosynthetic supply and metabolic demand of NADPH is important for operating stable CO_2_ fixation and avoiding oxidative stress by redox imbalance state^8,54^. In this study, it is suggested that accumulation of glycolytic intermediates acts as a buffering system that compensates for the gap between supply and demand of NADPH, and ensures rapid photosynthesis start. Indeed, lower accumulation of the glycolytic intermediates resulted in inefficient photosynthetic activity (Fig. 5). Previous study showed that 3PG is the most abundant in CBB cycle intermediates in three cyanobacterial species including *Synechocystis*^25^. This metabolite distribution is characteristic compared with that of heterotrophic organisms^12^ (Supplementary Fig. 11). Moreover, rapid consumption of phosphoglycerate upon illumination like Fig. 2 was also observed in algae, the chloroplast of spinach protoplasts, and leaves of wheat^55–57^. These facts support functional significance and conservation of accumulation of the glycolytic intermediates for robust photosynthesis.

## Methods

### Growth conditions

This work employed a glucose-tolerant wild-type strain of *Synechocystis*^58^ that was grown and stored on solid BG-11 media plates containing 1.5% Bacto agar. Pre-culturing was performed in 200 mL flasks containing 50 mL each of liquid BG-11 medium buffered with 20 mM 2-[4-(2-hydroxyethyl)-1-piperazinyl]ethanesulfonic acid (HEPES, pH 7.5); each flask was inoculated with cells harvested from the agar plates. These flasks then were cultured at 30 ºC in ambient air with shaking at 100 rpm; illumination at an intensity of 30 μmol m^−2^s^−1^ was provided by white light-emitting diode (LED) lights. The main culture was initiated by adding pre-culture grown for 4 days into 70 mL of the BG-11 medium to give an optical density of 0.05 at 730 nm (OD_730_). The growth condition of the main culture was the same as that of the pre-culture. The dry cell weight (DCW) was measured after harvesting cells by centrifugation and lyophilization. The obtained relationship between the OD_730_ value and DCW (DCW = 0.1998 × OD_730_ × V + 0.0974; R^2^=0.982; n=9; V is volume of cell suspension) was used for estimating DCW from the OD_730_ value.

### Preparation of ^13^C isotope-labeled internal standard (^13^C-IS)

*Synechocystis* cells in pre-culture were inoculated to an OD_730_ of 0.05 in a serum bottle containing 60 mL modified BG-11 medium supplemented with 100 mM HEPES (pH 7.5) and 25 mM NaH^13^CO_3_ (in place of the unlabeled Na_2_CO_3_ included in the standard BG-11 medium). The culture serum bottle was sealed, followed by flushing with N_2_ gas to remove unlabeled ambient CO_2_. The growth conditions used for the labeled culture were the same as those used for the pre-culture and main culture. Cells grown for 3 days were harvested and re-suspended in fresh modified BG-11 to avoid pH increase during cultivation with NaH^13^CO_3_ as the sole carbon source. The labeled culture then was grown under the same conditions for another 3 days, followed by extraction of uniformly labeled metabolites, as described below.

The entire volume of the labeled culture was harvested by centrifugation, washed with 1 mM NaH^13^CO_3_ solution, and then re-suspended in 9 mL of 1 mM NaH^13^CO_3_. The resulting cell suspension was illuminated for 90 s with white LED light at an intensity of 200 μmol m^−2^s^−1^ (to maximize extraction yield of CBB cycle intermediates), and placed into a mixture of 9 mL PCI (pH 7.9, Nacalai Tesque, Kyoto, Japan) and 60 μL 0.5 M N-cyclohexyl-3-aminopropanesulfonic acid (CAPS, pH 10.6) for metabolite extraction. The extraction mixture was centrifuged at 4 ºC at 8,000 ×*g* for 3 min. Twelve to 13 fractions of the water layer were then aliquoted into microtubes and lyophilized.

One of the dried aliquots was dissolved in 20 μL of standard mixture containing prepared unlabeled compounds. The sample was analyzed by capillary electrophoresis-mass spectrometry (CE-MS) (CE, Agilent G7100; MS, Agilent G6224AA LC/MSD TOF; Agilent Technologies, Palo Alto, CA, USA) as described previously^59^. Absolute concentrations of the uniformly labeled isotopomer (*C*_*n*_) were derived as follows:

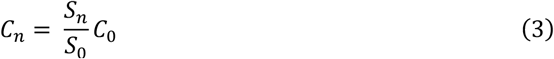

where, *S*_*n*_ and *S*_*0*_ are the peak areas of uniformly ^13^C-labeled (namely, the number of ^13^C atoms in a *n*-carbon compound is equal to *n*) and unlabeled standard signals, respectively, and *C*_*0*_ is the prepared concentration of the unlabeled standard.

### Metabolite extraction with rapid sampling and analysis

To perform metabolite sampling for analyses of absolute concentration and ^13^C incorporation after the transition from dark to light, a main culture grown for 6 days was dark-adapted for 12 h. A calculated volume (90/OD_730_, in mL) of the dark-adapted cell culture was harvested by centrifugation at 8000×*g* for 5 min. The cells were washed with 4 mL of 1 mM NaHCO_3_ (for absolute concentration analysis) or 1 mM NaH^13^CO_3_ (for ^13^C incorporation analysis), and harvested again by centrifugation. The washed cells were resuspended in 6.8 mL of 1 mM NaHCO_3_ (for absolute concentration analysis) or 1 mM NaH^13^CO_3_ (for ^13^C incorporation analysis). Nine aliquots (750 μL each) of the resulting cell suspension were then loaded into clear pipet tips (Supplementary Fig. 12). Up to this point, light irradiation was avoided during the procedure to keep the cells in the dark-adapted state. Irradiation of all the aliquots with white LED light at an intensity of 200±10 μmol m^−2^s^−1^ or 30±2 μmol m^−2^s^−1^ was initiated, followed by sampling of the cells at the indicated time points by adding into 750 μL PCI supplemented with ^13^C-IS dissolved in 20 μL water (for absolute concentration analysis) or with 5 μL of 0.5 M CAPS (pH 10.6) (for ^13^C incorporation analysis). The cell suspension before illumination was extracted as the nominal 0-s sample. To precisely determine the amount of extracted cells, the OD_730_ of the final cell suspension was measured. After the extraction mixtures were centrifuged at 4 ºC at 12,000×*g* for 2 min, harvested aliquots of the water layer were lyophilized.

The lyophilized samples were dissolved in 20 μL of 0.4 mM piperazine-1,4-bis(2-ethanesulfonic acid) (PIPES) and analyzed by CE-MS as described above. Absolute concentrations of each metabolite (*C’*) were derived as follows:

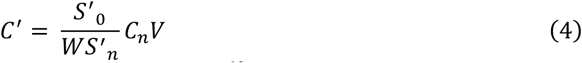

where, *S’*_*0*_ and *S’*_*n*_ are the peak areas of unlabeled and uniformly ^13^C-labeled IS signals, respectively, *W* represents DCW, and *V* is the sample volume for CE-MS analysis. The intracellular concentration (mM) was calculated by using the values: 5 × 10^7^ cells/mL/OD_730_ ^60^ and 4.4 × 10^− 15^ L/cell^61^. ^13^C fractions were calculated as described previously^30^.

### Short-term dynamic metabolic flux analysis (STD-MFA)

To estimate the temporal changes of 18 fluxes in the metabolic map shown in Supplementary Fig. 8, 18 metabolite accumulation rates *r*_*i*_(t) in equation (1) were derived from the slope taken between two consecutive concentration data points shown in Fig. 2. Accumulation rates of metabolites under the detection limit (E4P, R5P, GAP <0.001 nmol/mg DCW) and peak-unidentified metabolites (BPG and SBP) were set to zero. In addition, the CO_2_ fixation rate derived from the slope taken between two consecutive data points shown in Fig. 3 was regarded as the metabolic flux of RuBisCO (i.e., the conversion of RuBP to 3PG). The set of 19 equations, including the 18 unknown fluxes in the network, was represented in matrix notation as follows:

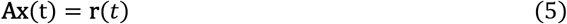

where, A is the 19 × 18 matrix of stoichiometric coefficients, **x**(t) is the 18-dimensional flux vector, and r(t) is the 19-dimensional metabolite accumulation rate vector. The specific form of equation (4) is shown in Supplementary Fig. 8b. The weighted least squares solution to equation (5) is as follows:

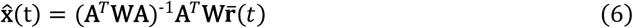

where, the superscripts *^* and ^−^ denote estimated and measured quantities (respectively), and **W** is a 19-dimensional weighting diagonal matrix, whose diagonal elements are the reciprocal of the variance of the measured metabolic accumulation rate values. Variance values for the accumulation rate of E4P, R5P, GAP, BPG and SBP, and for the CO_2_ fixation rate, were set to 10^−15^ (i.e., a value that was effectively zero). To evaluate the reliability of the estimated fluxes, the metabolite accumulation rates were calculated as follows:

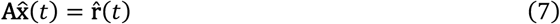

The estimated and measured metabolite accumulation rate values are compared in Supplementary Fig. 9.

## Acknowledgements

We thank Prof. C. Miyake (Kobe University) and Dr. Ginga Shimakawa (Kwansei Gakuin University) for kindly providing the mutants used in this work. This work was supported by the Mirai Program [grant number JPMJMI19E4] from the Japan Science and Technology Agency (JST) of the Ministry of Education, Culture, Sports, Science, and Technology (MEXT), Japan, and JSPS KAKENHI Grant Number 21J00113 and 22K15142.

## Author contributions

K.T. and T.H. designed the research. K.T. and M.M. performed research. K.T. and T.S. analyzed data. K.T., T.S., C.J.V., A.K. and T.H. wrote the article.

