## Supplementary information for "Dark accumulation of downstream glycolytic intermediates confers robust initiation of photosynthesis in cyanobacteria"

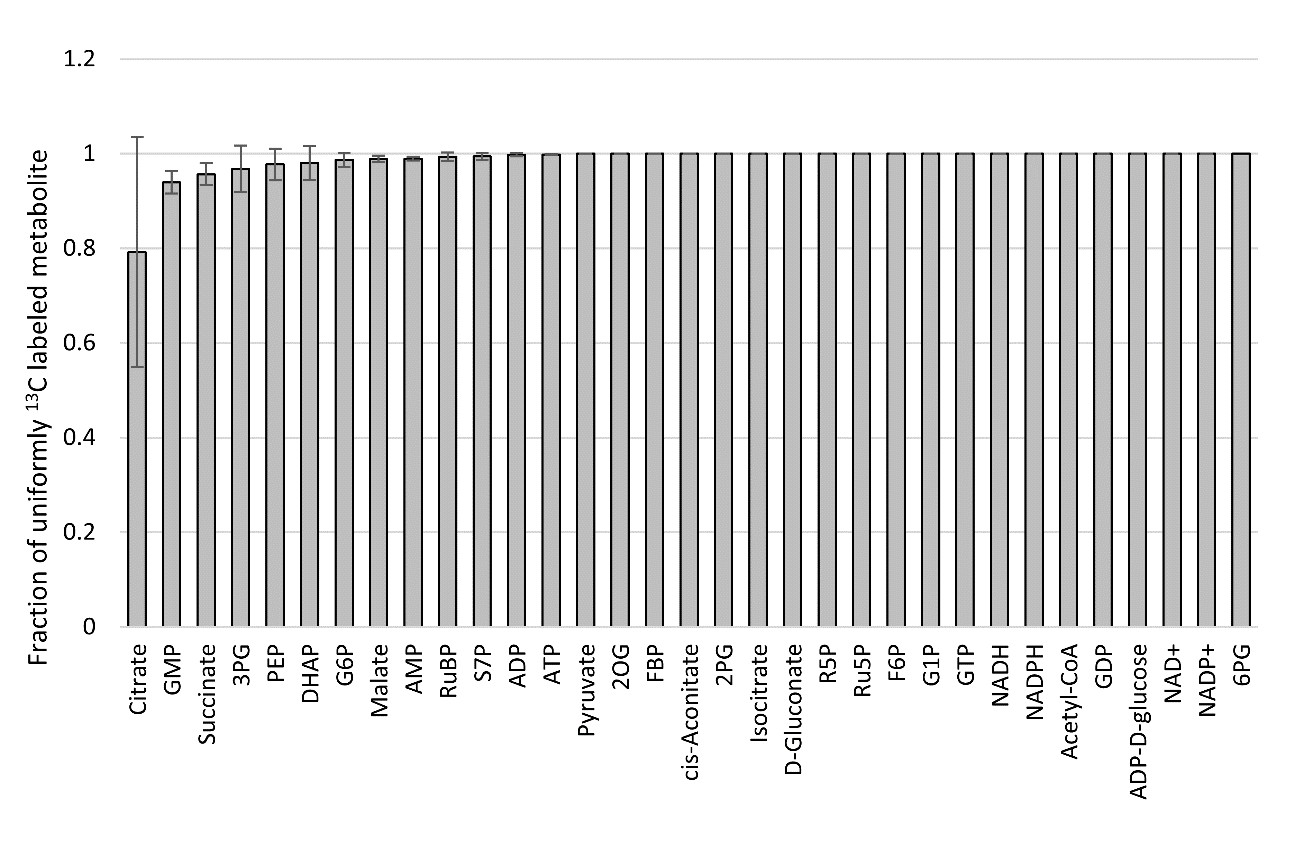

**Supplementary Fig. 1 Fraction of uniformly labeled metabolite in ^13^C internal standards.** Values are calculated as the ratio of uniformly labeled peak area to sum of peak areas of uniformly labeled and unlabeled signals. Values are the mean ± SD (bars) of six different ^13^C-internal standards. Abbreviations are the same as in the Figure 2 legend.

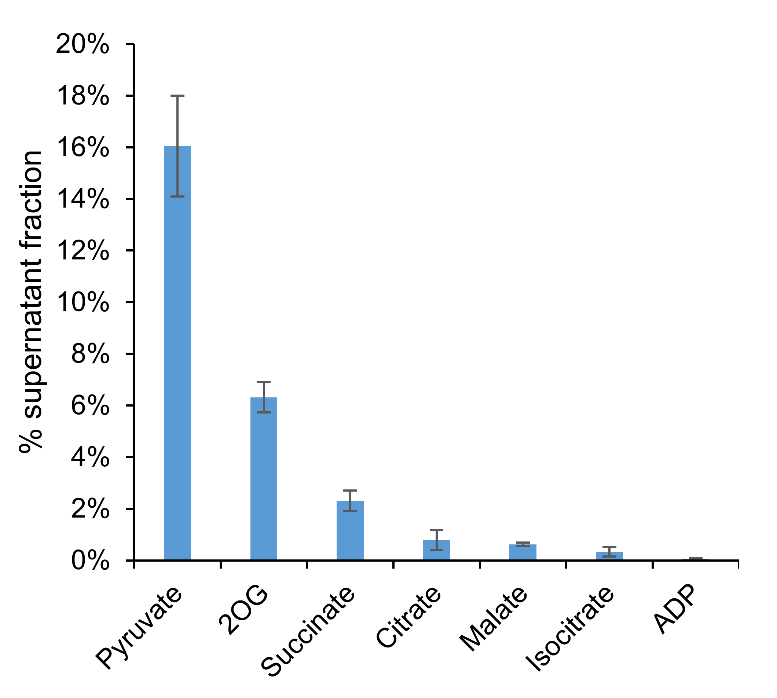

**Supplementary Fig. 2 Fractions of extracellular metabolites at 2 min after start of illumination with an intensity of 200 μmol m^−2^ s^−1^.** Cell suspensions in 1 mM NaHCO_3_ are prepared in the same manner as used for absolute concentration measurement. After illumination for 2 min, cell suspensions are centrifuged, followed by harvesting supernatant as the extracellular fraction. After lyophilization, these extracellular metabolite samples are analyzed using capillary electrophoresis-mass spectrometry (CE-MS). The extracellular fractions are determined from the ratio of peak area of extracellular samples to that of the extracted samples. Values are the mean ± SD (bars) of three biological replicates. ADP, Adenosine diphosphate; 2OG, 2-oxoglutarate.

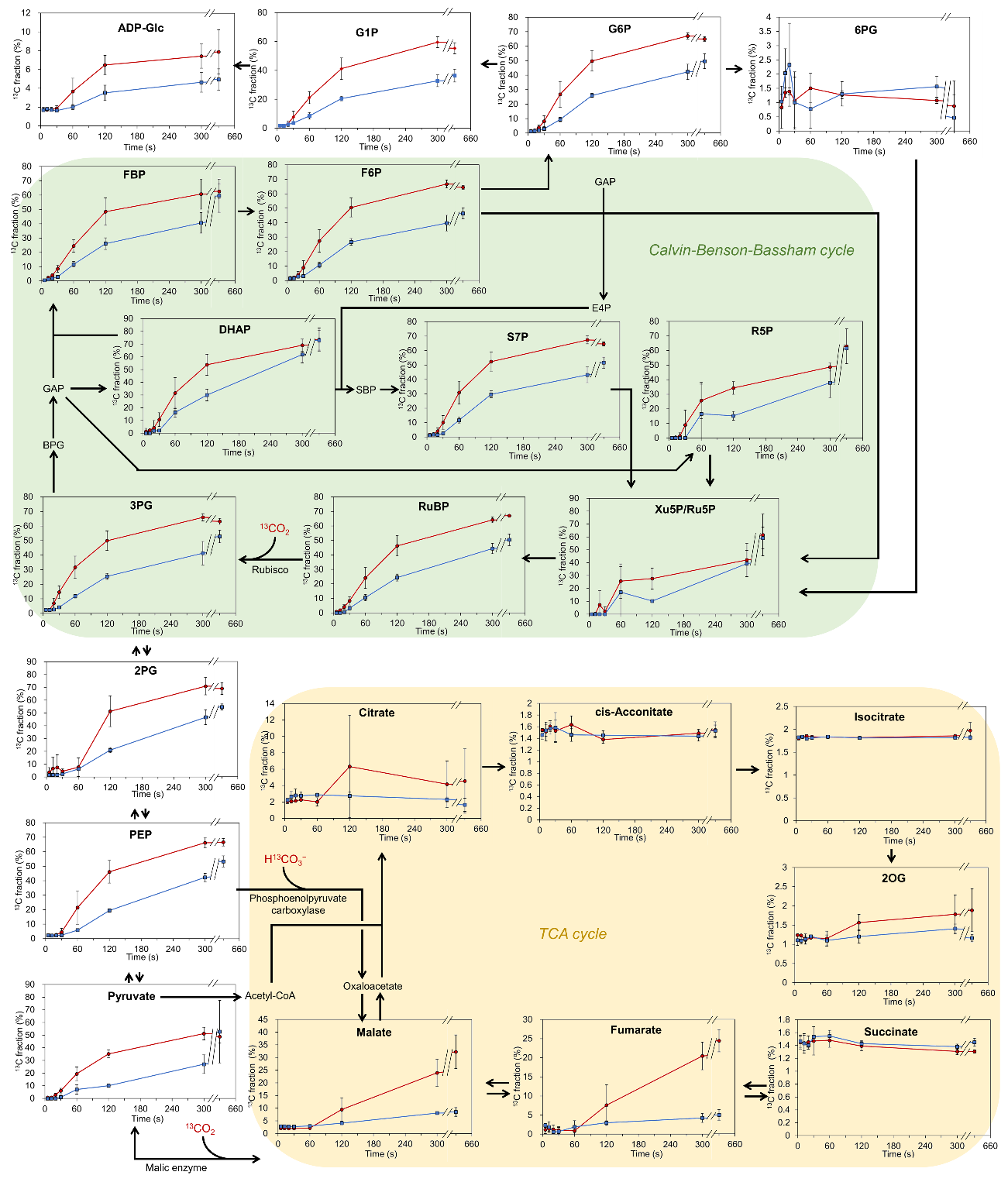

**Supplementary Fig. 3 Time course ^13^C metabolite fractions following light irradiation at 200 μmol m^−2^ s^−1^ (red line) or 30 μmol m^−2^ s^−1^ (blue line).** Values are the mean ± SD (bars) of three biological replicates. Black arrows indicate metabolic pathways. Abbreviations are the same as in the Figure 2 legend.

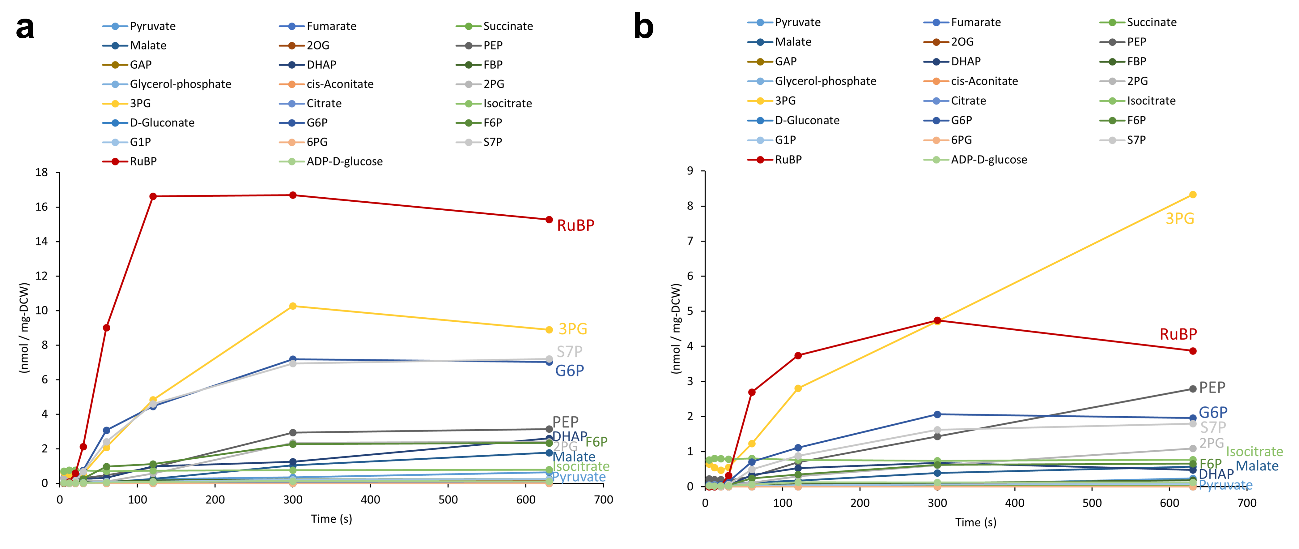

**Supplementary Fig. 4 Time course of ^13^C level in each metabolite.** Number of ^13^C atoms is calculated as the mean of ^13^C enrichment multiplied by mean metabolite concentration and the number of C atoms in the molecule for the two light intensities: (a) 200 μmol m^−2^ s^−1^, (b) 30 μmol m^−2^ s^−1^. Abbreviations are the same as in the Figure 2 legend.

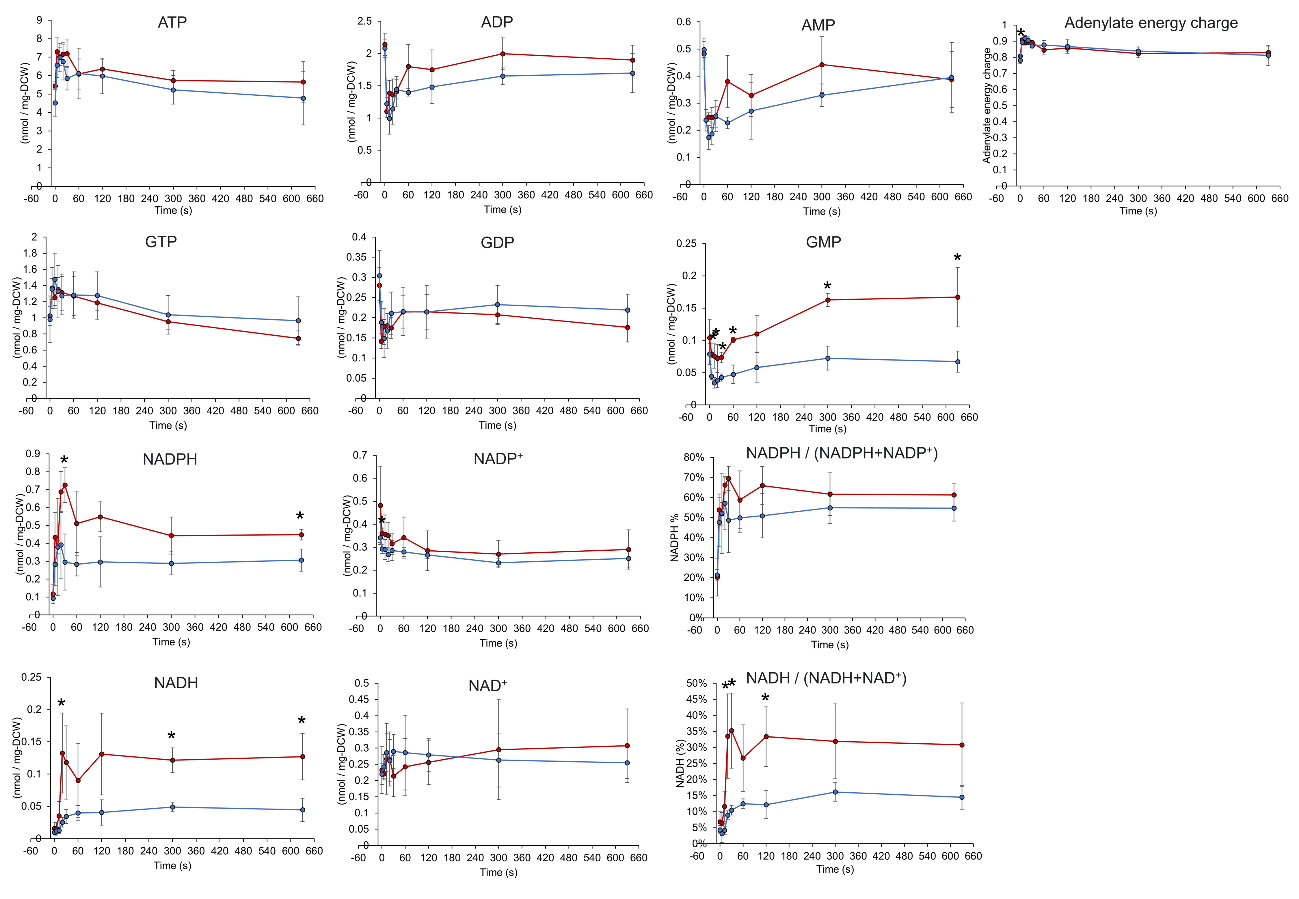

**Supplementary Fig. 5 Concentration transients of cofactors during photosynthetic induction.** Intracellular metabolites are extracted at the indicated times after transition from darkness to light-irradiated conditions at the intensity of 200 μmol m^−2^ s^−1^ (red line) or 30 μmol m^−2^ s^−1^ (blue line). Values are the mean ± SD (bars) of three independent experiments. Significant differences between the two light intensities were evaluated by a two-tailed non-paired Student’s t test (* *p* < 0.05). Abbreviations: ADP, Adenosine diphosphate; AMP, adenosine monophosphate; ATP, adenosine triphosphate; GDP, Guanosine diphosphate; GMP, Guanosine monophosphate; GTP, Guanosine triphosphate; NAD(P), nicotinamide adenine dinucleotide (phosphate).

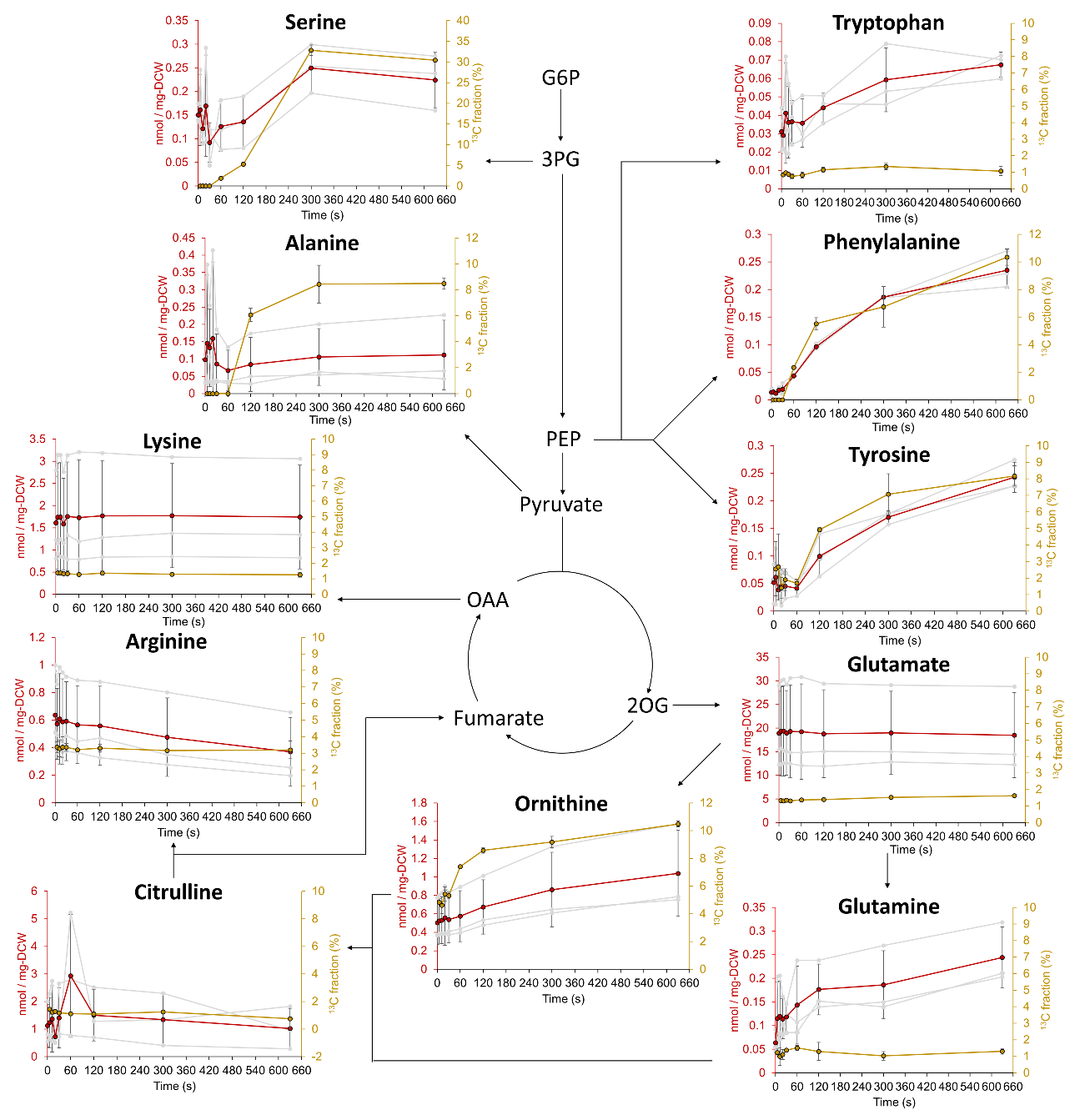

**Supplementary Fig. 6 Concentration and ^13^C fraction transients of amino acids during photosynthetic induction of cells illuminated at** **an intensity of 200 μmol m^−2^ s^−1^.** Intracellular metabolites are extracted at the indicated times after transition from darkness to light-irradiated conditions at an intensity of 200 μmol m^−2^ s^−1^. Red lines and yellow lines indicate metabolite concentration and ^13^C fraction, respectively. Simplified central metabolic pathways are indicated by arrows. Values of concentrations and ^13^C fractions are the mean ± SD (bars) of three and two independent experiments, respectively. Values of metabolite concentrations for individual experiments are indicated by gray lines. Abbreviations are the same as in the Figure 2 legend.

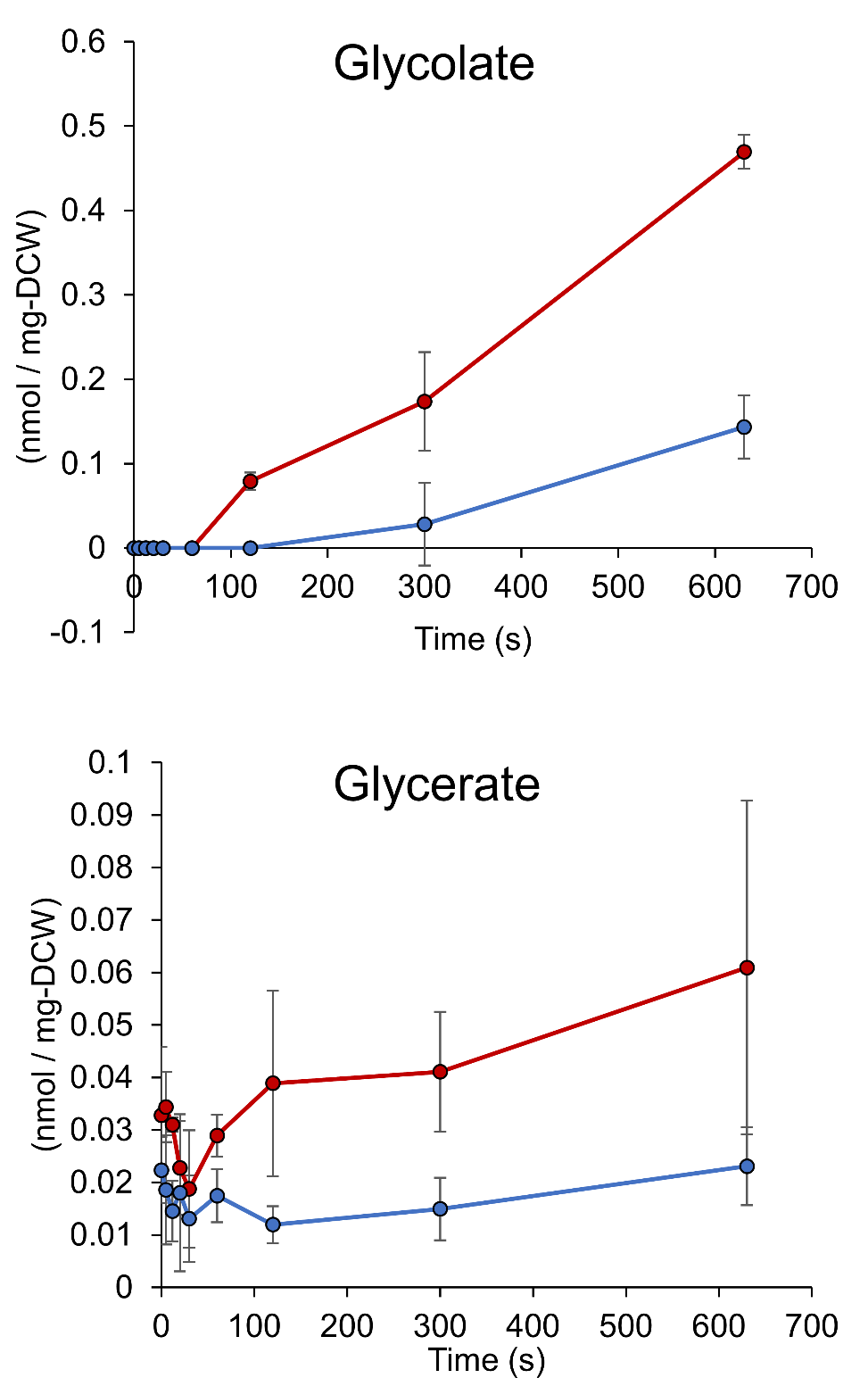

**Supplementary Fig. 7 Concentration transients of intermediates in photorespiration pathway during photosynthetic induction.** Intracellular metabolites are extracted at the indicated time after transition from darkness to light-irradiated conditions at intensities of 200 μmol m^−2^ s^−1^ (red line) or 30 μmol m^−2^ s^−1^ (blue line). Values are the mean ± SD (bars) of three independent experiments. DCW, dry cell weight.

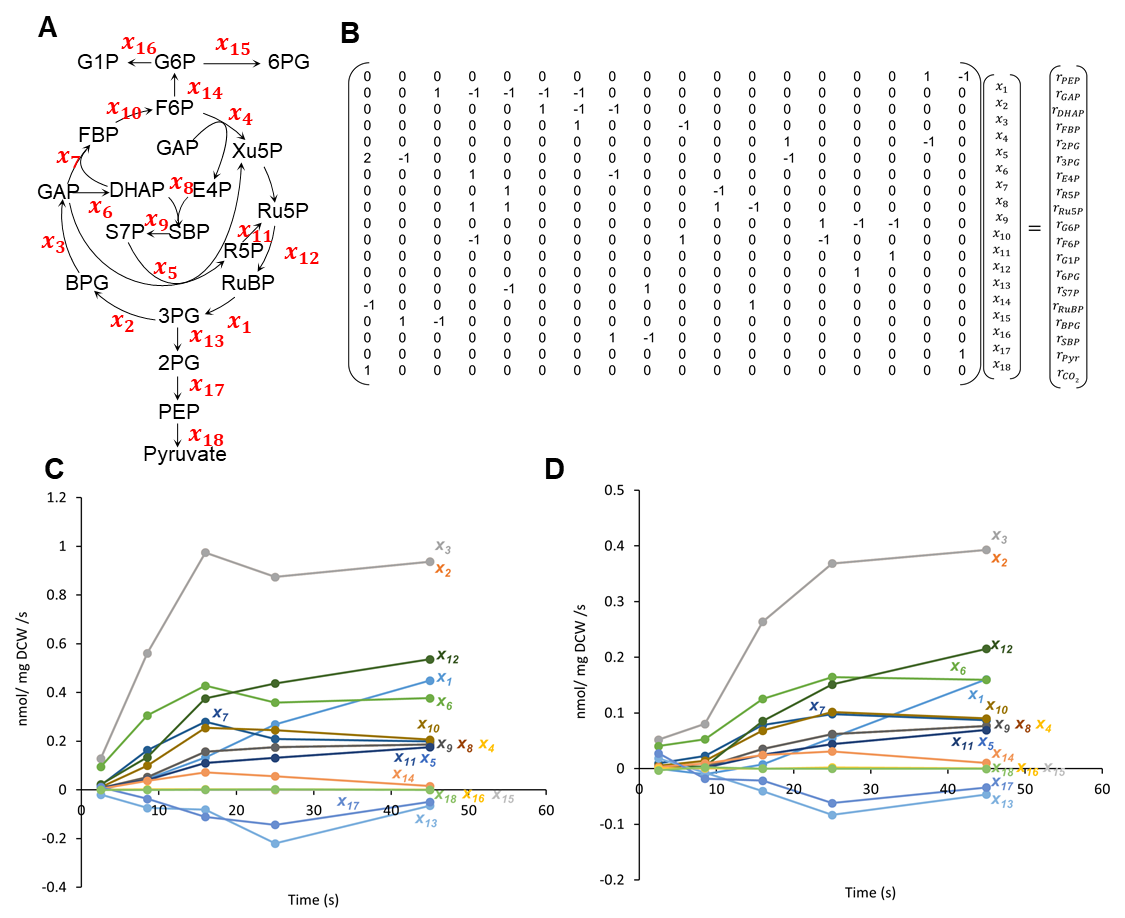

**Supplementary Fig. 8 Short-term dynamic metabolic flux analysis (STD-MFA).** (a) Metabolic pathways included in STD-MFA calculations. The experimental data of absolute concentration and ^13^C incorporation change allow us to simplify the metabolic model during photosynthetic induction. $x_{i}$ represents net flux of *i*th metabolic pathway. (b) Detailed form of the mass balance equation used for STD-MFA. For more details, see Methods section. (c, d) Transients of estimated metabolic fluxes during photosynthetic induction in illuminated conditions at (c) 200 μmol m^−2^ s^−1^ or (d) 30 μmol m^−2^ s^−1^. Since the fluxes are estimated using metabolite accumulation rates derived from two consecutive concentration data points, flux values are plotted at a time point corresponding to the average (mean) of the two time points. Abbreviations are the same as in the Figure 2 legend.

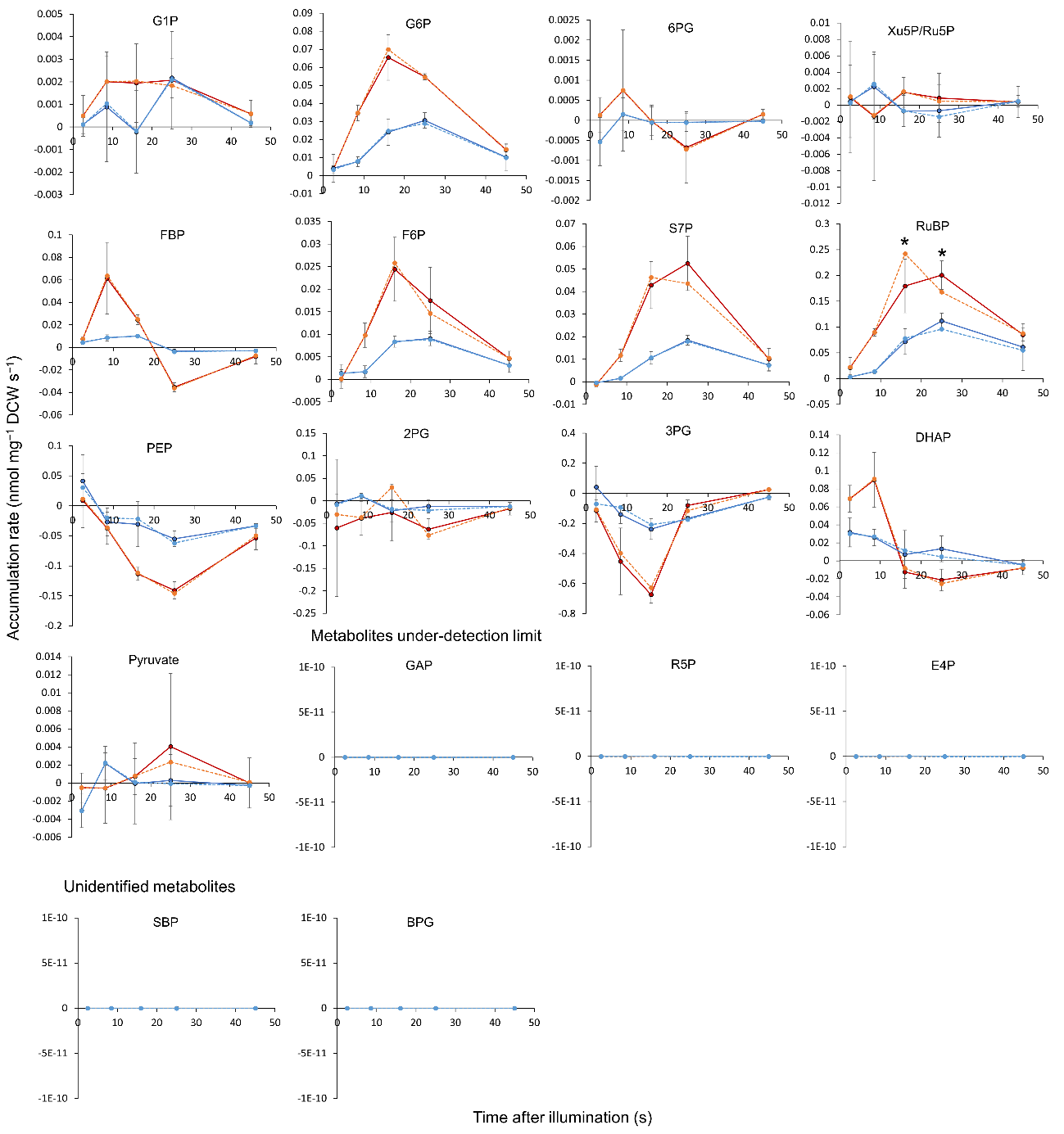

**Supplementary Fig. 9 Measured and estimated metabolic accumulation rates.** Measured accumulation rates at light intensities of 200 μmol m^−2^ s^−1^ (red solid line) or 30 μmol m^−2^ s^−1^ (blue solid line) are derived from slope between two consecutive concentration data points shown in Fig. 2. Estimated accumulation rates at light intensities of 200 μmol m^−2^ s^−1^ (orange dotted line) or 30 μmol m^−2^ s^−1^ (blue dotted line) are shown for validation of the short-term dynamic metabolic flux analysis (STD-MFA) results. Black arrows indicate metabolic pathways. Abbreviations are the same as in the Figure 2 legend. Values are plotted at the average (mean) time point of the two time points used for the concentration data points. Values of measured accumulation rates are the mean ± SD (bars) of three independent experiments. Asterisks indicate that the estimated value is out of range of the measured mean ± SD.

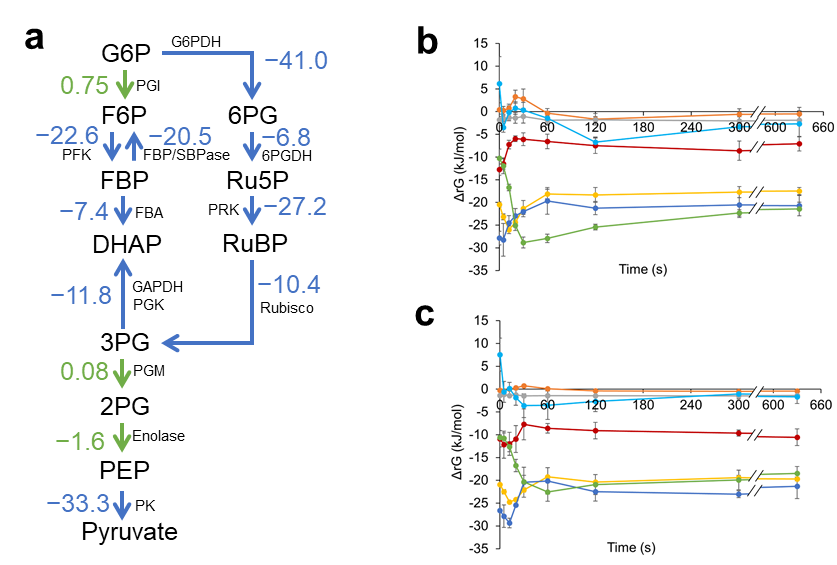

**Supplementary Fig. 10 Free energy analysis.** (a) Reaction Gibbs free energy change (Δ*_r_G*) values of glycolysis and pentose phosphate pathway in dark conditions are calculated from the absolute concentrations, and shown in the metabolic map. Blue and green arrows indicate Δ*_r_G* is below −5, between −5 and 5, respectively. (b,c) Time course of Δ*_r_G* at the illuminated light intensities of 200 μmol m^−2^ s^−1^ (b) and 30 μmol m^−2^ s^−1^ (c). Δ*_r_G* values of reactions (3PG to DHAP: red line, 3PG to 2PG: orange line, FBP to F6P: yellow line, Ru5P to RuBP: blue line, RuBP to 3PG: green line, 2PG to PEP: gray line, DHAP to FBP: light blue) were derived based on equation (2). Values are the mean ± SD (bars) derived from three independent experiments.

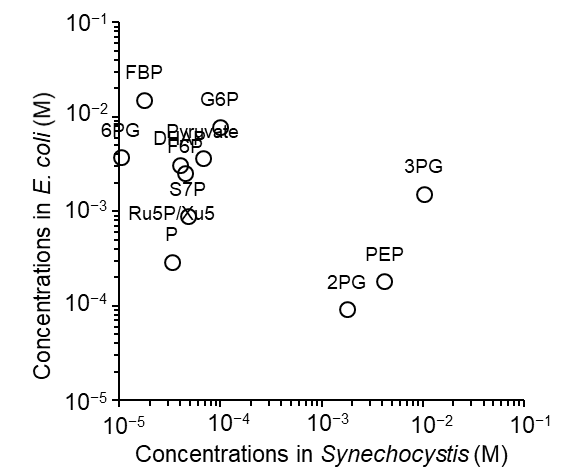

**Supplementary Fig. 11 Comparison of absolute metabolite concentrations between *Escherichia coli* and *Synechocystis*.** Metabolite concentrations of *E. coli* are derived from Park et al^12^. Metabolite concentrations of *Synechocystis* are the mean of six dark samples (Fig. 2, t = 0).

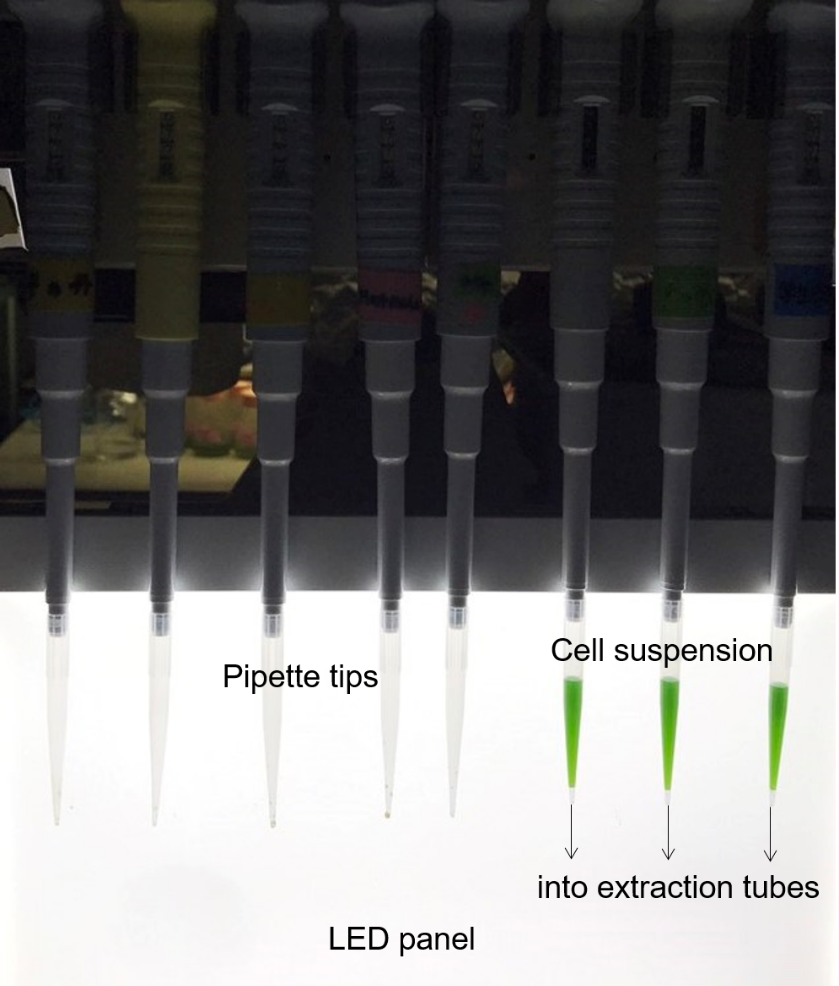

**Supplementary Fig. 12 Rapid sampling system used in this study.** Setting the cell suspensions in pipette tips prior to light irradiation permits rapid sampling during photosynthetic induction into extraction tubes containing phenol-chloroform-isoamyl alcohol (PCI). LED, Light emitting diode.

**Supplementary Table 1 standard reaction Gibbs energy change from eQuilibrator database.**

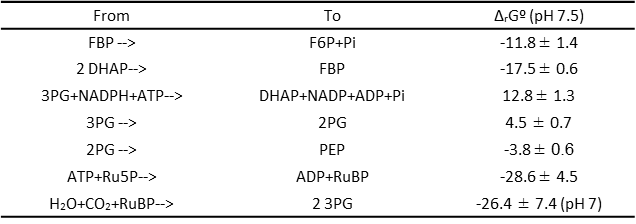
